# Expression of a single inhibitory member of the Ly49 receptor family is sufficient to license NK cells for effector functions

**DOI:** 10.1101/2024.06.04.597367

**Authors:** Sytse J. Piersma, Shasha Li, Pamela Wong, Michael D. Bern, Jennifer Poursine-Laurent, Liping Yang, Diana L. Beckman, Bijal A. Parikh, Wayne M. Yokoyama

## Abstract

Natural killer (NK) cells recognize target cells through germline-encoded activation and inhibitory receptors enabling effective immunity against viruses and cancer. The Ly49 receptor family in the mouse and killer immunoglobin-like receptor family in humans play a central role in NK cell immunity through recognition of MHC class I and related molecules. Functionally, these receptor families are involved in licensing and rejection of MHC-I-deficient cells through missing-self. The Ly49 family is highly polymorphic, making it challenging to detail the contributions of individual Ly49 receptors to NK cell function. Herein, we showed mice lacking expression of all Ly49s were unable to reject missing-self target cells *in vivo*, were defective in NK cell licensing, and displayed lower KLRG1 on the surface of NK cells. Expression of Ly49A alone on a H-2D^d^ background restored missing-self target cell rejection, NK cell licensing, and NK cell KLRG1 expression. Thus, a single inhibitory Ly49 receptor is sufficient to license NK cells and mediate missing-self *in vivo*.

## Introduction

Natural killer (NK) cells are innate lymphoid cells (ILCs) that can mediate effective immunity against viruses and cancer through direct lysis and cytokine production (Huntington et al., 2020; Piersma and Brizić, 2021). NK cells recognize their target cells through integration of signals by germline-encoded activation and inhibitory receptors (Long et al., 2013). These inhibitory receptors include members of the Ly49 family in the mouse and killer immunoglobin-like receptor (KIR) family in humans and they prevent killing of healthy cells through recognition of MHC class I (MHC-I) (Colonna and Samaridis, 1995; Karlhofer et al., 1992). Host cells may lose surface MHC-I expression in response to virus infection or malignant transformation. As a result, these cells become invisible to CD8^+^ T cells, but simultaneously become targets for NK cells through “missing-self” recognition (Kärre et al., 1986).

The inhibitory Ly49 molecules appear to be responsible for missing-self recognition in mice (Babić et al., 2010; Belanger et al., 2012; Gamache et al., 2019; Parikh et al., 2020; Zhang et al., 2019). However, it has been sometimes challenging to draw definitive conclusions because the Ly49 family is highly polymorphic and differs between mouse strains. Moreover, multiple inhibitory Ly49 receptors within a single host can recognize a given MHC-I molecule while others apparently have no ligands and instead recognize other MHC-I alleles (Schenkel et al., 2013). Yet, the Ly49s for non-host MHC-I alleles are still expressed. For example, in the C57BL/6 background, Ly49C and Ly49I can recognize H-2^b^ MHC-I molecules that include H-2K^b^ and H-2D^b^, while Ly49A and Ly49G cannot recognize H-2^b^ molecules and instead they recognize H-2^d^ alleles. Still these Ly49s are expressed in C57BL/6 mice, so their individual contributions to missing-self rejection are unclear. Ly49A has also been implicated in recognition of the non-classical MHC-I molecule H2-M3, which is upregulated in response to exposure to N-formylated peptides (Andrews et al., 2012; Chiu et al., 1999). Importantly, the specificities of several Ly49s have been clearly established while others remain to be confirmed. For example, the binding of Ly49A to H-2D^d^ has been confirmed by crystallographic studies (Tormo et al., 1999), and validated by mutational analysis of both Ly49A and H-2D^d^ (Matsumoto et al., 2001; Wang et al., 2001). By contrast, the MHC-I specificities of other Ly49s have been primarily studied with MHC tetramers containing human β_2_-microglobulin (B2m), which is not recognized by Ly49A (Mitsuki et al., 2004), on cells overexpressing Ly49s (Hanke et al., 1999). Thus, the contributions of individual Ly49 receptors to NK cell effector function are confounded by expression of multiple receptors, some of which may be irrelevant to a given self-MHC haplotype, and multiple Ly49 alleles whose specificities are less well defined.

In addition to effector function in missing-self, Ly49 receptors that recognize their cognate MHC-I ligands are involved in licensing or education of NK cells to acquire functional competence. NK cell licensing is characterized by potent effector functions including IFNγ production and degranulation in response to activation receptor stimulation (Elliott et al., 2010; Kim et al., 2005). Like missing-self recognition, inhibitory Ly49s require SHP-1 for NK cell licensing which interacts with the ITIM-motif encoded in the cytosolic tail of inhibitory Ly49s (Bern et al., 2017; Kim et al., 2005; Viant et al., 2014). Moreover, lower expression of SHP-1, particularly within the immunological synapse, is associated with licensed NK cells (Schmied et al., 2023; Wu et al., 2021). Thus, inhibitory Ly49s have a second function that licenses NK cells to self-MHC-I thereby generating functionally competent NK cells but it has not been possible to exclude contributions from other co-expressed Ly49s.

The complex nature of the Ly49 family confounds our fundamental understanding of these receptors, particularly regarding their function *in vivo*. To better understand Ly49 function *in vivo*, several groups made mutant mouse lines with altered expression of the *Ly49* locus. This was pioneered by the Makrigiannis group, which targeted the *Ly49o* promoter in 129 ES cells and resulting mice were subsequently backcrossed to C57BL/6 background (Belanger et al., 2012). These mice were defective in rejection of MHC-I deficient target cells *in vivo* and exhibited reduced tumor control as well (Tu et al., 2014). However, there was limited surface expression of NKG2A as well as Ly49s so these target defects were not solely dependent on absence of Ly49s. Moreover, these mice likely carried 129 alleles of other genes in the NK gene complex (NKC) that are expressed on NK cells, display allelic polymorphisms and are genetically linked to *Ly49*. Following the development of mouse CRISPR engineering, the Dong group deleted the entire 1.4 Mb *Ly49* locus in the C57BL/6 NKC and showed that Ly49-deficient mice were unable to reject MHC-I deficient cells *in vivo* at steady state (Zhang et al., 2019). Besides loss of Ly49 expression, surface expression of other receptors, including KLRG1, NKG2A, NKG2D and CD94 were also reduced in these mice. Around the same time, we generated a mouse that contains a 66 Kb and 149 Kb deletion in the C57BL/6 Ly49 locus, resulting in loss of 4 Ly49 molecules including Ly49A and Ly49G (Parikh et al., 2020). The resulting ΔLy49-1 mice were also deficient in H2D^d^-restricted control of murine cytomegalovirus (MCMV), which was rescued by knock-in of *Ly49a* into the *Ncr1* locus. Thus, available genetic evidence suggest inhibitory Ly49 receptors are essential for licensing and missing-self recognition.

The current data, however, do not take into account that individual Ly49s are stochastically expressed on NK cells, and multiple receptors are simultaneously expressed on individual NK cells, resulting in a diverse Ly49 repertoire of potential specificities on overlapping subsets of NK cells (Dorfman and Raulet, 1998; Kubota et al., 1999; Smith et al., 2000). As a result, not all NK cells express a specific Ly49 receptor and most NK cells express multiple Ly49 molecules, making it difficult to study the biology of a specific Ly49 receptor without genetic approaches. However, *Ly49* genes are highly related and clustered together, resulting in a high concentration of repetitive elements (Makrigiannis et al., 2005), complicating the capacity to target individual *Ly49* genes for definitive analysis. Moreover, in ΔLy49-1 mice which had an intact *Ly49d* coding sequence, the percentage of NK cells expressing Ly49D was markedly reduced, even though Ly49D was otherwise expressed at normal levels, suggesting a regulatory locus control region within the deleted fragments (Parikh et al., 2020). Such data raise the possibility that the large genetic deletions of the Ly49 locus may affect other NK cell receptors in the NKC that contribute to NK cell function. Thus, expression of individual Ly49 receptors without confounding effects of other Ly49s and potentially other NKC genes is needed to validate the conclusions from study of mice lacking Ly49 expression.

Here we studied the role of an individual Ly49 receptor in NK cell function. To this end, we deleted all NK cell Ly49 genes using CRISPR/Cas9 and confirmed the role of the *Ly49* family in missing self and licensing. Subsequently, we expressed Ly49A in isolation under control of the *Ncr1* locus in Ly49-deficient mice on a H-2D^d^ background to show that a single inhibitory Ly49 receptor expressed by all NK cells is sufficient for licensing and mediating missing-self *in vivo*.

## Results

### NK cells from CRISPR-generated mice lacking all NK cell-related Ly49 molecules display reduced KLRG1 expression

To investigate the role of individual Ly49 molecules, we generated a mouse that lacked all expressed Ly49 receptors. We targeted the remaining *Ly49* region in our previously published ΔLy49-1 mouse (Parikh et al., 2020) with guide RNAs targeting *Ly49i* and *Ly49q*. The resulting mouse contained a fusion between *Ly49i* and *Ly49q* with a deletion of the start codon and insertion of a fusion sequence that did contain a potential start site for a putative 5 amino acid polypeptide (Figure 1A). We confirmed that the 3’ deletion reported in the ΔLy49-1 mouse was unaffected, resulting in a frameshift and a premature stop codon after 9 amino acids in the fused *Ly49a/g* gene. Thus, genetic and sequencing analysis revealed all Ly49 genes were disrupted and we termed this mouse line Ly49KO (Figure 1A).

**Figure 1.**
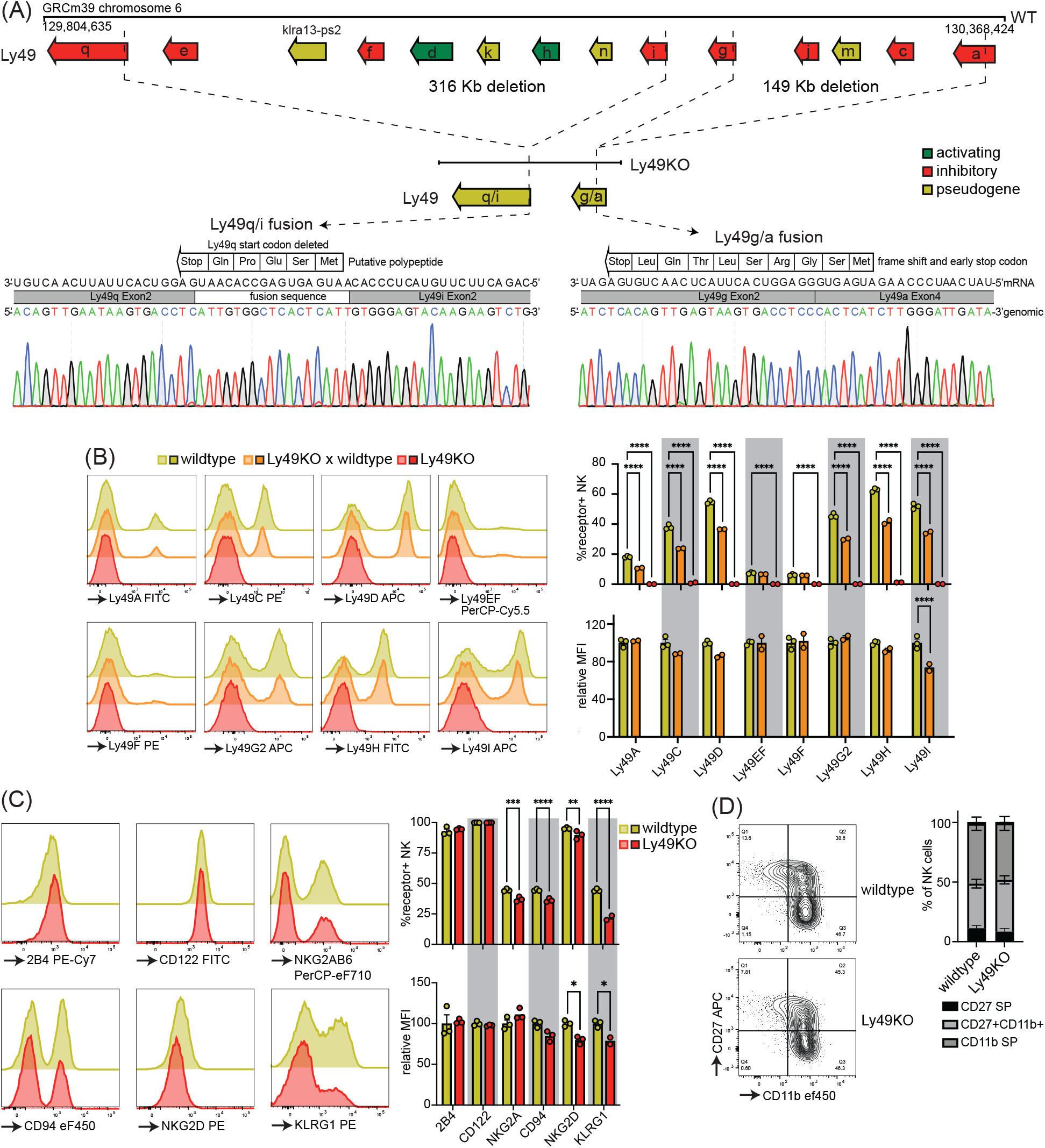
Mice generated to lack all NK-related Ly49 molecules using CRISPR have NK cells that display alterations in select surface molecules. (**A**) Genetic map of the Ly49 locus of wildtype C57BL/6 and Ly49KO mice and Sanger sequencing of the fusion sequences in the Ly49KO mice. (**B**) Ly49 receptor expression on splenic NK cells of the indicated mice. (**C**) Surface receptor expression on splenic NK cells from indicated mice. (**D**) Expression of the maturation markers CD27 and CD11b on splenic NK cells from indicated mice. MFI, median fluorescent intensity. Error bars indicate SEM; ns, not significant; \**p* < 0.05, \*\**p* < 0.01, \*\*\**p* < 0.001, and \*\*\*\**p* < 0.0001.

Flow cytometry confirmed loss of cell surface expression of Ly49 molecules in homozygous Ly49KO mice (Figure 1B). NKG2A, CD94, and NKG2D molecules that are encoded by the NKG2 locus, located next to the Ly49 locus, were still expressed, albeit at marginally lower frequencies (Figure 1C) (Yokoyama and Plougastel, 2003). Unrelated molecules 2B4 and CD122 were unaffected. In heterozygous Ly49KO mice, the percentages of NK cells expressing Ly49A, Ly49C, Ly49D, Ly49G2, Ly49H, and Ly49I were reduced by 33-41%. The median fluorescent intensity (MFI) for Ly49I was reduced by 26% in Ly49I^+^ NK cells in heterozygous Ly49KO mice, while the other Ly49s did not display significant differences in MFI. Consistent with apparent dependence of normal KLRG1 expression on MHC-I expression (Corral et al., 2000) and previous reports (Zhang et al., 2019), Ly49KO NK cells showed a 51% reduction in KLRG1 expression (Figure 1C). NK cells in Ly49KO mice displayed similar maturation to wildtype NK cells, based on expression of the surface markers CD27 and CD11b (Figure 1D). Thus, Ly49KO mice specifically lack all Ly49 molecules and display moderate alterations in select surface molecules while showing otherwise normal numbers of apparently mature NK cells.

### Ly49-deficient NK cells are defective in licensing and rejection of MHC-I deficient target cells

Inhibitory Ly49-positive NK cells can be licensed through recognition of cognate MHC-I molecules, resulting in a phenotype of increased IFNγ production following plate-bound anti-NK1.1 stimulation (Kim et al., 2005). NKG2A has been implicated in NK cell licensing by the non-classical MHC-I molecule Qa1 (Anfossi et al., 2006), to eliminate potential confounding effects by this interaction, effector functions of NKG2A-NK cells were evaluated as described before (Bern et al., 2017). Stimulation of Ly49KO NK cells anti-NK1.1 resulted in a 73% reduction in IFNγ production as compared to wildtype NK cells, similar to unlicensed NK cells from H-2K^b^ and H-2D^b^ double deficient (KODO) mice (Figure 2A). Both Ly49KO and KODO NK cells produced high amounts of IFNγ in response to phorbol 12-myristate 13-acetate (PMA) plus ionomycin (Figure 2B), indicating that their IFNγ production machinery is intact. Thus, these results confirm that Ly49 molecules are required for the NK cell licensed phenotype.

**Figure 2.**
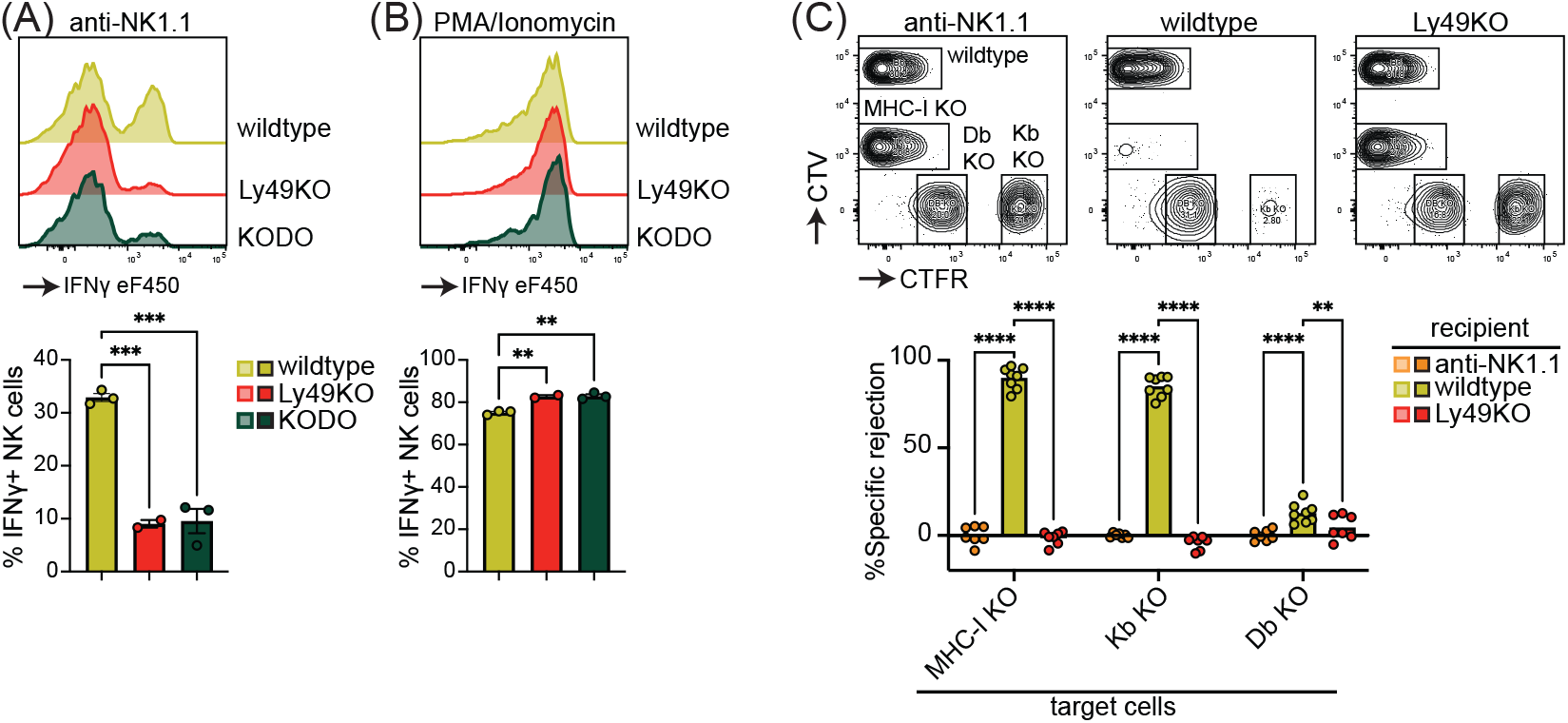
NK cell licensing and rejection of MHC-I deficient target cells is defective in Ly49KO mice. Splenocytes from the indicated mice were stimulated with plate bound anti-NK1.1 (**A**) or PMA/ionomycin (**B**) and IFNγ production by NKG2A-NK cells and analyzed by flow cytometry. (**C**) *In vivo* cytotoxicity assay against H-2K^b^, H-2D^b^ and full MHC-I deficient targets. Splenocytes from WT, H-2Kb, H-2Db and MHC-I deficient mice were differentially labeled with CTV and CTFR as indicated. Mixture of labeled target cells were injected i.v. into wildtype, Ly49KO, and anti-NK1.1-depleted mice. Target cells were analyzed in spleens by flow cytometry 2 days after challenge. KODO, H-2K^b^ x H-2D^b^ knock out; MHC-I KO, KODO x B2m knockout; MFI, median fluorescent intensity. Error bars indicate SEM; ns, not significant; \**p* < 0.05, \*\**p* < 0.01, \*\*\**p* < 0.001, and \*\*\*\**p* < 0.0001.

To investigate the capability of Ly49KO NK cells to reject MHC-I deficient target cells, we challenged anti-NK1.1 NK cell-depleted, wildtype, and Ly49KO mice with a mixture of wildtype, B2m deficient x KODO (MHC-I KO), H-2D^b^ KO, and H-2K^b^ KO splenocytes that were differentially labeled with CellTrace Far Red (CTFR) and CellTrace Violet (CTV) (Figure 2C). While wildtype mice efficiently rejected MHC-I KO and H-2K^b^ KO target cells, only 13% of H-2D^b^ KO target cells were rejected by wildtype mice, indicating that missing-self recognition in the C57BL/6 background depends on the absence of H-2K^b^ rather than H-2D^b^. None of the target cell populations were rejected in the Ly49KO mice, comparable to wild type controls depleted of NK cells. Thus, Ly49 molecules mediate NK cell-dependent MHC-I deficient target cell killing *in vivo* under steady-state conditions and rejection of MHC-I deficient target cells is predominantly controlled by H-2K^b^ in the H-2^b^ background.

### Expression of Ly49A in Ly49-deficient H-2D^d^ transgenic mice rescues KLRG1 expression

To investigate the potential of a single inhibitory Ly49 receptor on mediating NK cell licensing and missing-self rejection, the Ly49KO mice were backcrossed to H-2D^d^ transgenic KODO (D8-KODO) Ly49A KI mice that express *Klra1* cDNA encoding the inhibitory Ly49A receptor in the *Ncr1* locus encoding NKp46 and its cognate ligand H-2D^d^ but not any other classical MHC-I molecules (Parikh et al., 2020). Ly49A expression in the resulting Ly49KO/Ly49A KI D8-KODO mice closely follows NKp46 expression because NKp46^-^ NK1.1^+^ NK cells in the bone marrow of these mice do not express Ly49A, while virtually all the NKp46^+^ NK1.1^+^ NK cells in the bone marrow and spleen express Ly49A (Figure 3A). Ly49KO/Ly49A KI D8-KODO NK cells expressed robust levels of Ly49A, albeit at lower MFI as compared to Ly49A expression on D8-KODO NK cells, consistent with prior observations with wild type Ly49A and H2D^d^ (Held et al., 1996; Karlhofer et al., 1994). NK cells were able to fully mature in Ly49KO D8-KODO and Ly49KO/Ly49A KI D8-KODO mice as we observed similar percentage of mature CD27^-^ CD11b^+^ NK cells in spleen, bone marrow, and liver (Figure 3B). While there was a modest significant increase in immature CD27^+^ CD11b^-^ NK cells in the bone marrow of Ly49KO D8-KODO and Ly49KO/Ly49A KI D8-KODO mice, no differences were observed in NK cell maturation in spleen and liver. The decrease in the frequency of KLRG1^+^ NK cells observed in the Ly49KO NK cells on the H-2^b^ background was recapitulated in the D8-KODO background (Figure 3C). Intriguingly, expression of Ly49A in Ly49KO/Ly49A KI D8-KODO mice rescued KLRG1 expression and resulted in similar levels of KLRG1 as D8-KODO NK cells. Taken together, Ly49A engineered to be encoded within the NKp46 locus was efficiently expressed as an isolated Ly49 receptor and supported KLRG1 expression.

**Figure 3.**
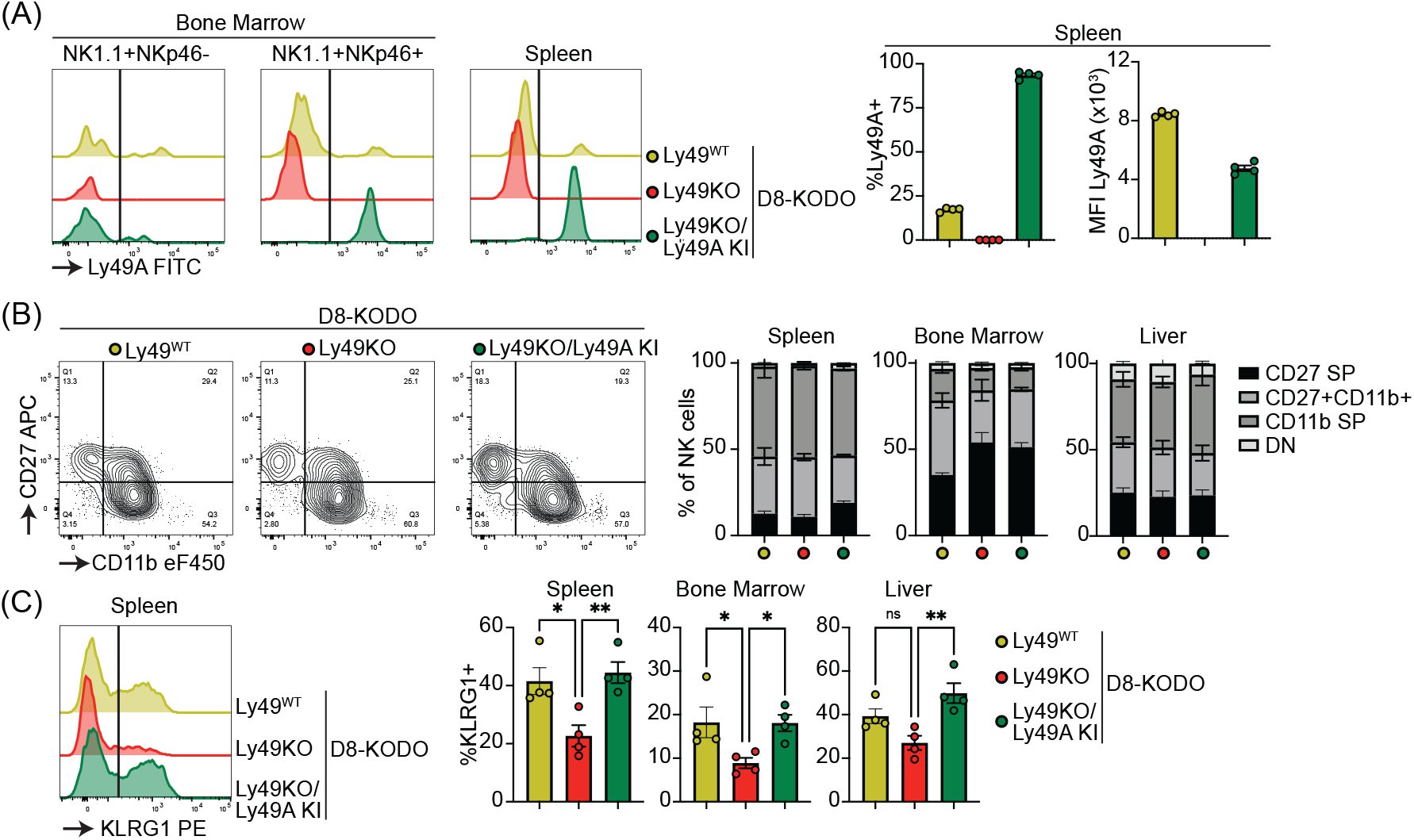
Ly49A is efficiently expressed in Ncr1-Ly49A knockin mice and rescues KLRG1 expression in NK cells. Flow cytometric analysis of NK cells in D8-KODO, Ly49KO D8-KODO, and Ly49KO/Ly49A KI D8-KODO mice (**A**) Ly49A expression in NKp46^+^ and NKp46^-^ NK1.1^+^ NK cells in bone marrow and spleen of the indicated mice. (**B**) Expression of the maturation markers CD27 and CD11b on NK cells in spleen, bone marrow, and liver of indicated mice. (**C**) KLRG1 expression by NK cells in spleen, bone marrow, and liver of indicated mice. MFI, median fluorescent intensity. Error bars indicate SEM; ns, not significant; \**p* < 0.05, \*\**p* < 0.01, \*\*\**p* < 0.001, and \*\*\*\**p* < 0.0001.

### NK cells expressing Ly49A in isolation are fully licensed and capable of rejecting MHC-I deficient target cells

Next, we investigated the potential of Ly49A expression alone to mediate NK cell licensing and rejection of MHC-I deficient target cells. In D8-KODO mice, Ly49A^+^ NK cells displayed increased levels of IFNγ production and degranulation measured by CD107 in response to plate bound anti-NK1.1 stimulation as compared to all NK cells including unlicensed cells (Figure 4A). Similar to Ly49KO NK cells on the H-2^b^ background, Ly49KO NK cells on the D8-KODO background showed a 72% decrease in IFNγ production, but also showed a 59% decrease in degranulation as compared to Ly49A^+^ NK cells in D8-KODO mice. This impaired IFNγ production and degranulation was reversed in Ly49A KI NK cells on the Ly49KO D8-KODO background, that showed similar IFNγ production and degranulation to Ly49A^+^ NK cells on the D8-KODO background, indicating that Ly49A KI NK cells are licensed. Importantly, there were no differences among all NK cell populations in IFNγ production and degranulation in response to PMA/Ionomycin (Figure 4B), indicating that the IFNγ production and degranulation machinery are not affected in any of the mouse strains. Thus, the Ly49KO/Ly49A KI D8-KODO NK cells displayed a fully licensed phenotype comparable to licensed Ly49A^+^ NK cells in D8-KODO mice.

**Figure 4.**
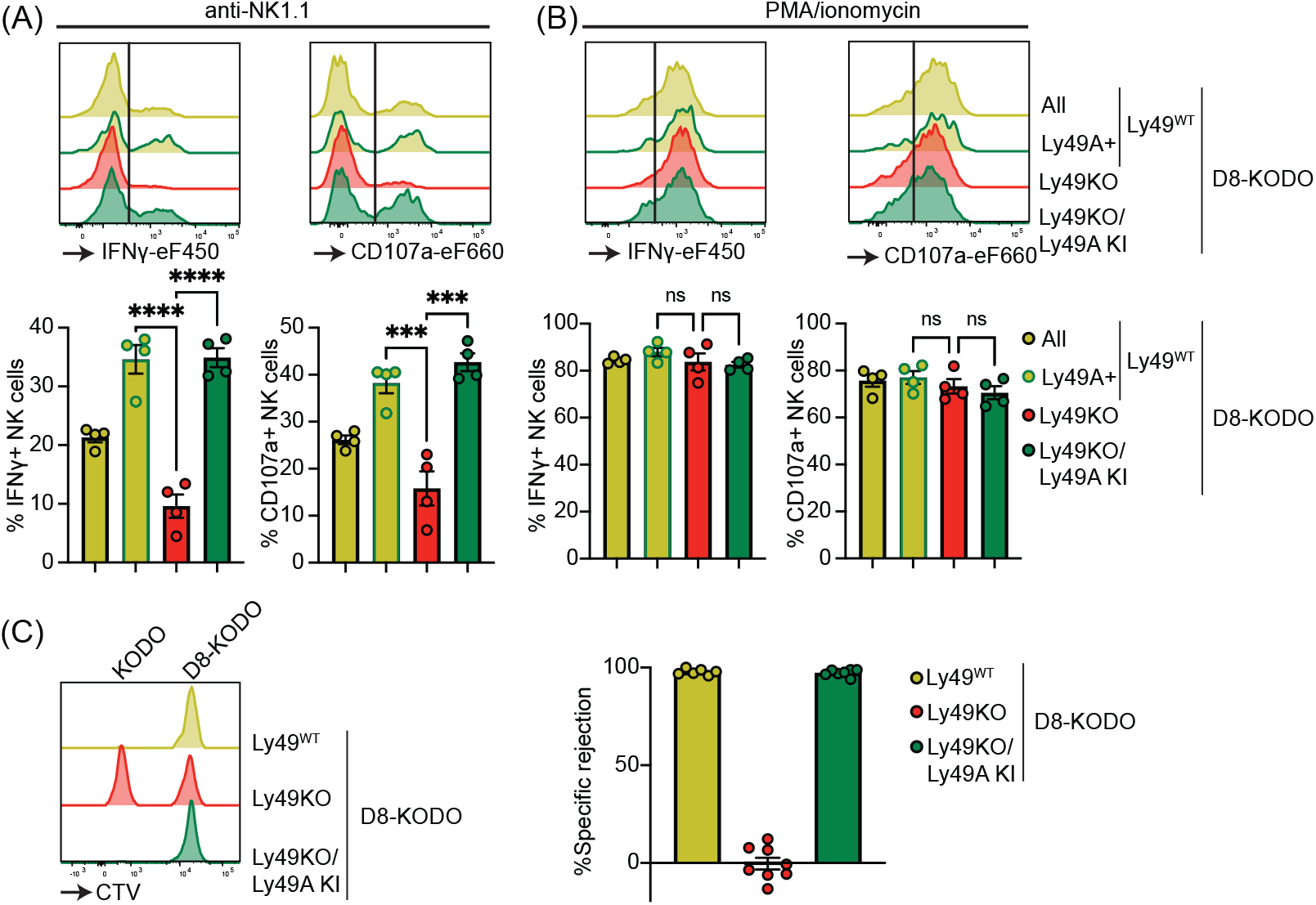
Expression of Ly49A in isolation is sufficient for NK cell licensing and missing-self rejection. Splenocytes from the indicated mice were stimulated with plate bound anti-NK1.1 (**A**) or PMA/ionomycin (**B**). IFNγ production and degranulation (CD107a) by NKG2A-NK cells were analyzed by flow cytometry. (**C**) Splenocytes from D8-KODO and KODO mice were differentially labeled with CTV as indicated. A mixture of labeled target cells were injected i.v. into D8-KODO, Ly49KO D8-KODO, and Ly49KO/Ly49A KI D8-KODO mice. Specific rejection of target cells was analyzed in spleens by flow cytometry 2 days after challenge. Error bars indicate SEM; ns, not significant; \*\*\**p* < 0.001, and \*\*\*\**p* < 0.0001.

Finally, we interrogated whether a single inhibitory Ly49 molecule would be sufficient to mediate missing-self rejection of MHC-I deficient cells. To this end D8-KODO, Ly49KO D8-KODO, and Ly49KO/Ly49A KI D8-KODO mice were challenged with a mixture of D8-KODO and KODO (MHC-I deficient) splenocytes that were differentially labelled with CTV. D8-KODO mice efficiently rejected KODO target splenocytes, while Ly49KO D8-KODO mice were unable to reject these cells. However, this defect was completely restored in the Ly49KO/Ly49A KI D8-KODO mice, demonstrating that a single inhibitory Ly49 receptor is sufficient to mediate missing-self rejection of cells lacking its MHC class I ligand.

## Discussion

Several mice with genetic modifications in the Ly49 complex have been developed to study the role of Ly49 receptors but they have limitations (Belanger et al., 2012; Bern et al., 2017; Gamache et al., 2019; Parikh et al., 2020; Zhang et al., 2019). A complicating factor in these studies is that multiple Ly49s, often with incompletely understood specificities, may be involved in target cell recognition. Here, we studied a mouse where a single Ly49 was under control of the *Ncr1* locus which is expressed on all NK cells on the background of a complete Ly49 KO. Consistent with previous reports (Belanger et al., 2012; Parikh et al., 2020; Zhang et al., 2019), Ly49-deficient NK cells were deficient in licensing and missing-self rejection, both on a H-2^b^ background and in the presence of a single classical MHC-I allele, H-2D^d^. While our results did not interrogate licensing by inhibitory receptors outside of the Ly49 receptor family, such as has been reported for NKG2A (Anfossi et al., 2006; Zhang et al., 2019), they do demonstrate that expression of Ly49A without other Ly49 family members can mediate NK cell licensing. Moreover, we found that Ly49 receptors are required and sufficient for missing-self rejection under steady-state conditions. However, these observations do not rule out involvement of other inhibitory receptors under specific inflammatory conditions. For example, NKG2A contributes to rejection of missing-self targets in poly(I:C)-treated mice (Zhang et al., 2019). Finally, expression of the inhibitory Ly49A in isolation did not alter NK cell numbers or maturation, indicating that the Ly49s do not affect these parameters of NK cells. Yet the NK cells were fully licensed in terms of IFNγ production and degranulation *in vitro* and efficiently rejected MHC-I deficient target cells *in vivo*. Thus, a single Ly49 receptor is capable to confer the licensed phenotype and missing-self rejection *in vitro* and *in vivo*.

We observed that rejection of H-2K^b^-deficient targets was more potent than H-2D^b^-deficient target splenocytes, comparable to previous observation<u>w</u>s (Johansson et al., 2005). These data indicate that H-2K^b^ is more efficiently recognized by Ly49s as compared to H-2D^b^. This is further supported by early studies using Ly49 transfectants binding to Con A blasts showing that Ly49C and Ly49I can bind to H-2D^b^-deficient but not H-2K^b^-deficient cells (Hanke et al., 1999), despite the caveat of testing binding to cells overexpressing Ly49s in these studies. Our studies indicate that the efficiency of Ly49-dependent missing-self rejection depends on characteristics of specific MHC-I alleles recognized by cognate Ly49 receptors *in vivo*.

KLRG1 is an inhibitory receptor that recognizes E-, N-, and R-cadherins to inhibit NK cell cytotoxicity (Ito et al., 2006). KLRG1 is expressed on a subset of mature NK cells and can be upregulated in response to proliferation in a host with lymphopenia (Huntington et al., 2007). Consistent with previously published results (Zhang et al., 2019), we observed decreased KLRG1 expression in Ly49-deficient NK cells. The *Ly49* gene family as well as *Klrg1* is located within the NKC on chromosome 6 (Yokoyama and Plougastel, 2003), thus an effect of regulatory elements deleted in Ly49KO mice cannot be excluded. This is further emphasized by studies of our Δ Ly49-1 mouse which expresses Ly49D on fewer NK cells (Parikh et al., 2020). The *Ly49d* gene appears intact and Ly49D^+^ NK cells expressed Ly49D at normal levels, suggesting the absence of a regulatory element in the deleted regions. Here, we showed expression of only Ly49A, encoded in the *Ncr1* locus located on chromosome 7, in Ly49KO mice on a H-2D^d^ background restored KLRG1 expression in NK cells from different tissues, indicating that inhibitory Ly49 receptors rather than regulatory elements influence KLRG1 expression. Moreover, NK cell KLRG1 expression is modulated by MHC-I molecules (Corral et al., 2000). Therefore, KLRG1 expression may be modulated as a consequence of NK cell licensing through inhibitory Ly49 receptors.

The Ly49 family is stochastically expressed on NK cells, resulting in a NK cell repertoire with different combinations of Ly49 receptors on individual NK cells. Not all Ly49 alleles are equally expressed which has been suggested to be dependent on allelic exclusion (Held et al., 1995). Epigenetic control including DNA-methylation, histone modification, and regulatory elements have been linked to Ly49 expression (Kissiov et al., 2022; McCullen et al., 2016; Rouhi et al., 2006; Saleh et al., 2004). We observed that in Ly49KO heterozygous mice the percentage of Ly49^+^ NK cells was reduced for each Ly49 molecule, yet the expression level measured by MFI was not affected except for Ly49I, indicating that loss of one allele does not affect Ly49 surface levels. Nonetheless, alternate mechanisms may control Ly49 expression as was observed for Ly49I. Expression of *Ly49a* within the *Ncr1* locus resulted in ubiquitous Ly49A expression in NK cells, albeit at lower levels compared to Ly49A^+^ D8-KODO NK cells. Despite the lower expression levels of Ly49A, Ly49A KI NK cells were fully licensed and efficiently eliminated MHC-I-deficient target cells, suggesting that minor alterations in expression levels of various Ly49s on individual NK cells may not affect NK cell functions. In conclusion, these data show that expression of a single inhibitory Ly49 receptor is necessary and sufficient to license NK cells and mediate missing self-rejection under steady state conditions *in vivo*.

## Materials and Methods

### Animals

C57BL/6 (stock # 556) mice were purchased from Charles Rivers laboratories, B2m-deficient (stock # 2087) were purchased from Jackson laboratories, H-2K^b^ x H-2D^b^ double-deficient (stock # 4215; KODO) mice were purchased from Taconic Farms. D8 is a H-2D^d^ transgenic mouse that has been previously described (Bieberich et al., 1986) and was provided by D. Marguiles, National Institute of Allergy and Infectious Diseases, Bethesda, MD. D8-KODO mice have been previously generated by crossing D8 transgenic mice to a KODO background (Choi et al., 2011). ΔLy49-1 and Ly49A KI mice were previously generated in our laboratory (Parikh et al., 2020). In Ly49A KI mice the stop codon of Ncr1 encoding NKp46 is replaced with a P2A peptide-cleavage site upstream of the Ly49A cDNA, while maintaining the 3’ untranslated region. All mice were maintained within the Washington University animal facility in accordance with institutional ethical guidelines under protocol number 21-0090. All experiments utilized sex- and age-matched mice.

### Generation of Ly49KO mice

The remaining Ly49 receptors in ΔLy49-1 mice were targeted using CRISPR/Cas9 as previously described (Parikh et al., 2015a). Briefly, the Ly49 locus was targeted with gRNAs directed against Ly49q (5’-ACCCATGATGAGTGAGC<u>AGG</u>-3’) and Ly49i (5’-TGAGACTTCATAAGTCTTCA<u>AGG</u>-3’), with the PAM sequence underlined. For the mRNA microinjections, 20ng of each guide and 100ng of Cas9 mRNA were used. Deletions in the Ly49 locus were screened using the primers 5’-GCCCATCTGGCTTCCTTTCT-3’ (*Ly49q*-Rv), 5’-CAAGCCCCGATGAGATGGAT-3’ (*Ly49i*-Rv), and GGATCAGTCCATGTCAGGGTT (*Ly49i*-Fw) yielding a 409 bp wildtype and a 552 bp mutant band and confirmed using Southern blot analysis (data not shown). To minimize off-target CRISPR/Cas9 effects, candidate founder mice were backcrossed to C57BL/6 mice for 2 generations then crossed to derive homozygous Ly49KO mice. Deletions were verified by Sanger sequencing (Azenta Life Sciences) in homozygous Ly49KO mice using the PCR primers *Ly49q*-Rv and *Ly49i*-Fw for the *Ly49q/i* fusion sequence and the primers AACCAAGCCCCAATGAGATC (*Ly49g*-Rv) and TGGGTCAGTCCATGTCAGTG (*Ly49a*-Fw) for the *Ly49a/g* fusion sequence resulting in 552bp and 409bp products, respectively.

### Flow cytometry

Fluorescent-labeled antibodies Ly49D (clone 4D11), Ly49EF (CM4), Ly49F (HBF-719), Ly49G2 (eBio4D11), Ly49H (3D10), Ly49I (YLI-90), 2B4 (eBio244F4), CD122 (TM-b1), NKG2AB6 (16a11), NKG2ACE (20D5), CD94 (18d3), NKG2D (CX5), CD27 (LG.7F9), CD11b (M1/70), IFNγ (XMG1.2), CD107a (eBio1D4B), NKp46 (29A1.4), CD3(145-2C11), CD4 (RM4-5), CD8 (53-6.7), TCRB (H597), and CD19 (eBio1D3) were purchased from Thermo Fisher Scientific; Ly49A (YE1/48.10.6), KLRG1 (2F1), and NK1.1 (PK136), were purchased from Biolegend; Ly49C (4LO311) was purchased from Leinco Technologies. Cells were stained with fixable viability dye eF506 (Thermo Fisher Scientific), continued by staining of cell surface molecules in 2.4G2 hybridoma supernatant to block Fc receptors. For intracellular staining, cells were fixed and stained intracellularly using the BD Cytofix/Cytoperm Fixation/Permeabilization Kit (BD Bioscience) according to manufacturer’s instructions. Samples were acquired using FACSCanto (BD Biosciences) and analyzed using FlowJo software (BD Biosciences). NK cells were defined as singlet Viability-NK1.1+NKp46+CD3-CD19- or Viability-NK1.1+NKp46+CD4-CD8-TCRB-CD19-.

### *In vitro* stimulation assays

Stimulation of splenic NK cells was performed as previously described (Parikh et al., 2020; Piersma et al., 2019). Briefly, 1 - 4 µg/ml anti-NK1.1 (clone PK136, Leinco Technologies) in PBS was coated in 24-well plates for 90 min at 37ºC. Plates were washed with PBS and 5 × 10^6^ splenocytes were added per well. In parallel, splenocytes were stimulated with 200 ng/ml Phorbol myristate acetate (PMA; Sigma-Aldrich) and 400 ng/ml Ionomycin (Sigma-Aldrich). After 30 min incubation at 37ºC, Monensin (Thermo Fisher Scientific) and fluorescently labelled anti-CD107a antibody were added, cultures were incubated for an additional 7 hours at 37ºC and subsequently analyzed by flow cytometry.

### *In vivo* killing assays

*In vivo* killing assays were performed as previously described (Parikh et al., 2015b). Briefly, target splenocytes were isolated from C57BL/6, MHC-I deficient (TKO), H-2K^b^-deficient, H-2D^b^-deficient, KODO and D8-KODO mice. Indicated target splenocytes were differentially labelled with CellTrace violet, and/or CellTrace far red (Thermo Fisher Scientific). Target splenocytes were additionally labeled with CFSE to identify transferred target splenocytes from host cells. Target cells were mixed at equal ratios for each target and 2 × 10^6^ splenocytes per target were injected i.v. into naïve hosts. Where indicated NK cells were depleted with 100µg anti-NK1.1 (Leinco technologies) 2 days before target cell injection. Two days after challenge splenocytes were harvested and analyzed by flow cytometry. Target cell rejection was calculated using the formula [(1−(Ratio(KO target/wildtype target)_sample_/Ratio(KO target/wildtype target)_control_))×100].

## Statistics

All experiments were performed at least twice, and representative examples are shown. Cumulative data for 2 independent experiments is shown for *in vivo* killing assays. Statistical analysis was performed with Prism (GraphPad software) using unpaired *t*-tests and two-way ANOVA with corrections for multiple testing. Error bars in figures represent the SEM. Statistical significance was indicated as follows: ****, *p* < 0.0001; ***, *p* < 0.001; **, *p* < 0.01; *, *p* < 0.05; ns, not significant.

## Acknowledgments

We thank J. Michael White (Transgenic, Knockout, and Micro-Injection Core at Washington University) for CRISPR-Cas9 injections. This work was supported by National Institutes of Health grants R01-AI129545 to W.M.Y.

## Notes

### Competing Interest Statement

The authors have declared no competing interest.

### Summary of Updates

This revision include updated clarifications of the introduction, results section, discussion, and material and methods.

